# Intrinsic differences in hamster and mouse macrophage biology correlate with susceptibility to *L. donovani* infection

**DOI:** 10.1101/2025.08.01.668127

**Authors:** Paul Jenkins, Anne Danckaert, Thomas Cokelaer, Nassim Mahtal, Pascale Pescher, Gerald F. Späth

## Abstract

The basis for differential susceptibility to *Leishmania* (*L.*) *donovani* infection observed in individuals remains poorly understood. Here we address this important open question comparing bone marrow-derived macrophages from susceptible hamsters (hamBMDMs) and resistant mice (mBMDMs) to identify intrinsic cellular features that may contribute to host-specific outcomes. We first optimized and validated an experimental protocol for generating hamBMDMs, which closely resemble classical mouse BMDMs in terms of morphology, marker gene expression and phagocytic activity. Comparative transcriptomic analysis uncovered rodent-specific, intrinsic differences in the expression of metabolic and immune-related pathways known to influence susceptibility to intracellular *Leishmania* infection. *In vitro* infection assays confirmed the microbicidal capacity of hamBMDMs, while also revealing their increased permissiveness to *Leishmania* proliferation. In conclusion, the combined use of murine and hamster macrophage systems provides a powerful platform to dissect the molecular mechanisms underlying *L. donovani* survival and host resistance. Our improved protocol allows for the generation of large quantities of functionally validated hamster macrophages, enabling systems-level investigations in this important rodent model that more accurately reflects human infection dynamics than mice. This addresses a major bottleneck in experimental infections with *L. donovani,* but also other clinically relevant pathogens, such as *Mycobacterium* spp. and SARS-CoV-2, for which hamsters have been used to model human infection.

## Introduction

Macrophages serve as key antigen-presenting cells (APCs), bridging innate and adaptive immunity by instructing the immune system to mount appropriate responses against a variety of insults such as injury, cancer or infection (1). Given their central role in host defense, it is not surprising that these innate immune cells are the frequent target for intracellular viral, bacterial or protozoan pathogens, which exploit macrophages as host cells and subvert their functions to promote microbial survival, often resulting in severe immunopathologies (2, 3). Among such pathogens, the protozoan parasite *Leishmania* (L.) is widely regarded as a model organism for intracellular infection. These parasites have evolved sophisticated strategies to survive and proliferate within macrophages by resisting their cytolytic mechanisms, hijacking their immune functions and causing chronic infection (4–6).

Following inoculation of insect-stage promastigote parasites by blood feeding sandflies into vertebrate hosts, *Leishmania* parasites are phagocytosed by macrophages, where they differentiate into mammalian-stage amastigotes inside fully acidified phagolysosomes. Amastigote proliferation and subversion of host cell innate immune functions ultimately cause different immunopathologies that are collectively known as leishmaniases (7), which range from self-healing cutaneous leishmaniasis (CL) and disfiguring mucocutaneous leishmaniasis (MCL), to potentially fatal visceral leishmaniasis (VL).

VL, primarily caused by *L. donovani* and *L. infantum*, remains a significant global health concern. In 2023, 11,922 new VL cases were reported to WHO, with Eastern Africa (73%), Brazil (12%), and the Indian subcontinent (6%) identified as major hotspots (WHO, 2023). Clinical manifestations of VL include weight loss, fever, hepatosplenomegaly and anemia. While *L. donovani* causes the most severe forms of VL, not all infections are productive and lead to disease. Asymptomatic VL is common in endemic regions (8) and serves as a reservoir, facilitating parasite transmission and complicating control efforts (9–11). Several risk factors have been associated with VL severity, including malnutrition (12), microbial coinfection (13, 14), autoimmune disease (15) and host genetics (16–19). Despite their significance in disease epidemiology, the host determinants and immune mechanisms determining symptomatic versus asymptomatic VL remain poorly understood.

One critical yet underexplored factor in shaping VL disease outcome is the role of *Leishmania*– macrophage interactions. Shortly after infection, *L. donovani* rapidly remodels the host macrophage transcriptome, establishing permissive intracellular conditions for persistent infection (20). While transcriptomic analyses of symptomatic and asymptomatic VL patients have identified specific immune signatures in chronically infected cells (17, 21, 22), how these differences arise from the initial *Leishmania*–macrophage interactions and how they may promote opposite infection outcomes remain to be elucidated.

Here we address these important open questions leveraging macrophages from rodent models that recapitulate the two opposite outcomes of the human VL spectrum, i.e. *L. donovani* resistant mice and highly susceptible Syrian hamsters that eventually succumb to experimental VL. Mice are the most widely used animal models for VL research, yet they do not reproduce the progressive visceral pathology observed in humans and either rapidly clear (strains C3H and 129/SV) or control parasite burden (strains C57BL/6 and BALB/c)(23–25). In contrast, the Syrian golden hamster (Mesocricetus auratus) develops disease manifestations that closely parallel those seen in human VL caused by *L. donovani* (26–28).

To gain initial insight into intrinsic factor underlying the distinct disease outcomes observed in these two rodent systems, we employed a comparative transcriptomics approach using bone marrow-derived macrophages (BMDMs) from mice and hamsters. Building on a previous report (29), we optimized and biologically validated a protocol for generating hamster BMDMs (hamBMDMs) that yielded *bona fide* macrophages, as confirmed by their morphology, phagocytic capacity, and marker gene expression. Bulk RNAseq analysis on hamster and mouse BMDMs revealed species-specific macrophage expression profiles, with hamBMDMs showing features consistent with an immunosuppressive, M2-like polarization state, which was associated with higher parasite burden in *in vitro* infection experiments. Our results correlate intrinsic differences in rodent-specific macrophage biology to varying degrees of susceptibility to visceral leishmaniasis (VL), and set the stage for future, in-depth analyses of infection-dependent expression changes underlying symptomatic and asymptomatic VL.

## Material and Methods

### Ethics statement

Work on animals was performed in compliance with French and European regulations on care and protection of laboratory animals (EC Directive 2010/63, French Law 2013-118, February 6th, 2013). All animal experiments were approved by the Ethics Committee and the Animal welfare body of Institut Pasteur and by the Ministère de l’Enseignement Supérieur, de la Recherche et de l’Innovation (project n°#19683).

### Animals

Four to ten weeks old female Syrian golden hamsters (*Mesocricetus auratus*) and C57BL/6 mice (*Mus musculus*) were purchased from Janvier Labs and all animals were hosted in A3 animal facility at the Institut Pasteur. All animals were handled under specific, pathogen-free conditions in biohazard level 3 animal facilities (A3) accredited by the French Ministry of Agriculture for performing experiments on live rodents (agreement A75-15-01).

### Mammalian cell and parasite culture media

For both mouse and hamster, bone marrow-derived and peritoneal macrophages were cultured in Dulbecco’s Modified Eagle Medium (DMEM) high glucose (PAN Biotech, P04-03588), supplemented with 15% Fetal Bovine Serum (FBS, Gibco), 10 mM HEPES (Sigma Aldrich), 50 µM 2-Mercaptoethanol (Gibco), 100 units/ml Penicillin and 100 µg/ml Streptomycin solution (Corning).

*Leishmania donovani* parasites (Ld1S2D, MHOM/SD/62/1S-CL2D) derived from hamster splenic amastigotes were cultured in M199 medium supplemented with 25 mM HEPES (Gibco), 10% FBS (Gibco), 10 µM Adenine (Sigma life science), 100 units/ml Penicillin and 100 μg/ml Streptomycin (Corning), 2 mM L-glutamine (Gibco), 10 µg/ml folic acid (Sigma life science), 13.7 µM Hemin (Sigma life science), 4.2 mM NaHCO3 (from 7.5% stock at R.P.Normapur), 8 µM 6-Biopterin (Sigma life science), RPMI 1640 vitamin solutions 1X (Sigma) at neutral (7.4) or acidic (5.5) pH. Cells were cultured at 26°C in closed-cap culture flasks.

### Peritoneal macrophage preparation

Peritoneal exudate cells from hamster and mouse peritoneal cavity were recovered after injection of 10-20 ml of cold PBS with 5% FBS. Peritoneal fluids were collected, placed in a 50 ml tube and centrifuged for 10 min at 300 x g and 4°C. Cells were counted and plated in 12 well plates at 2.5 x10^6^ cells/well for RNA preparation in DMEM complete medium and incubated overnight at 37°C, 7.5% CO2. Non-adherent cells were removed from the wells and the adherent cells were treated with 350 µl of a guanidine-based lysis buffer (LBP), containing a large amount of chaotropic ions to denature proteins and protect RNA integrity (NucleoSpin® RNA plus kit, Macherey Nagel). Cell lysates in LBP were passed through a 1 ml syringe/26G needle and stored at -80°C until RNA extraction using Macherey Nagel extraction kit.

### BHK-conditioned medium and lysates for RNA extraction

Baby hamster kidney (BHK) fibroblast cells were cultured in DMEM (Gibco) supplemented with 5% of FBS (Gibco) and 500 µM of Penicillin/Streptomycin solution (Corning) at 37°C and 7.5% CO2. For the preparation of conditioned medium, BHK cells at 80-90% confluence were collected after Trypsin/EDTA (Sigma Aldrich) treatment. Cells were expanded in 162 cm^2^ ventilated flasks containing 50 ml of complete DMEM medium. After 1 day at 100% confluence the supernatant was recovered by centrifugation at 300 x g and passed through a 0.22 µm filter.

For RNA extraction, BHK cells were seeded in Petri dishes. At 90-100% confluence, cells were washed once with pre-warmed PBS and scraped with 700 µl of LPB lysis buffer from Macherey Nagel NucleoSpin® RNA plus kit according to the manufacturer’s instructions. Cell lysates in LBP were passed through a 1 ml syringe/26G needle and stored at -80°C until RNA extraction.

### Bone marrow progenitor cell isolation and differentiation into bone marrow-derived macrophages

All the procedures for the recovery of hamster and mouse progenitor cells were performed on ice. Progenitor cells were recovered from femurs and tibias by flushing the bone marrow with a 5 ml syringe/25G needle using cold PBS (Gibco) containing 2% of FBS. The cell suspensions were filtered through 100 µm cell strainers (Corning) into 50 ml tubes, completed with cold PBS-A (2% FBS) and centrifuged at 300 x g, 10 min, 4°C. The supernatants were discarded and red blood cells were lysed for 10 min on ice in 1X Gey buffer (10 ml of 1X Gey buffer is necessary for two hamster bones or four mouse bones). After incubation with Gey buffer, tubes were adjusted to 50 ml with cold PBS-A and centrifuged at 300 x g for 10 min at 4C°. The pellets were resuspended in 1 ml of cold complete DMEM medium, cells were counted and adjusted at the required concentrations.

For the differentiation of hamster progenitor cells into bone marrow-derived macrophages, the complete DMEM medium was supplemented with either 100 µg/ml of recombinant human-CSF1 (rhCSF1, Proteintech), 30% of BHK-conditioned medium (CM) or 100 µg/ml of recombinant human-CSF1 with 10% of BHK-conditioned medium (Mix). For the differentiation of mouse progenitor cells into bone-marrow derived macrophages, the complete DMEM medium used was supplemented with 50 µg/ml of recombinant mouse-CSF1 (rmCSF1, ImmunoTools). The bone-marrow cell suspensions recovered from the bones were diluted to 2.5×10^6^ cells/ml in complete DMEM medium. For hamster cells, 3 ml of suspension were seeded in a hydrophilic 6-well tissue culture plate (Falcon). For mouse cells, 12 ml of cell suspension were seeded on a 100 mm hydrophilic Petri dish (Falcon). Both hamster and mouse cell cultures were then incubated overnight at 37C°, 7.5% CO2. The next day, non-adherent hamster cells were recovered with the culture medium and diluted in 9 ml of complete DMEM medium and plated on a hydrophobic Petri dish (untreated Greiner bio-one, 100 mm diameter). Mouse non-adherent cells were collected with the culture medium, the suspension was adjusted to 36 ml with complete DMEM medium containing rmCSF1 and divided onto 3 hydrophobic Petri dishes (untreated Greiner bio-one, 100 mm). After three days of culture, 3 ml (for hamster cells) or 4 ml (for murine cells) of fresh complete DMEM medium containing respectively rhCSF1 or rmCSF1 were added to each Petri dish. Following three additional days of culture, loosely adherent and non-adherent cells were flushed from the Petri dish using a 10 ml pipette and collected in a 50 ml tube on ice. The suspension was then centrifuged at 300 x g for 10 min at 4C° and resuspended in 1 ml of cold complete DMEM medium for counting (Figure 1B). Cells were plated overnight at different concentrations according to the experiment to perform (see sections below).

**Figure 1:**
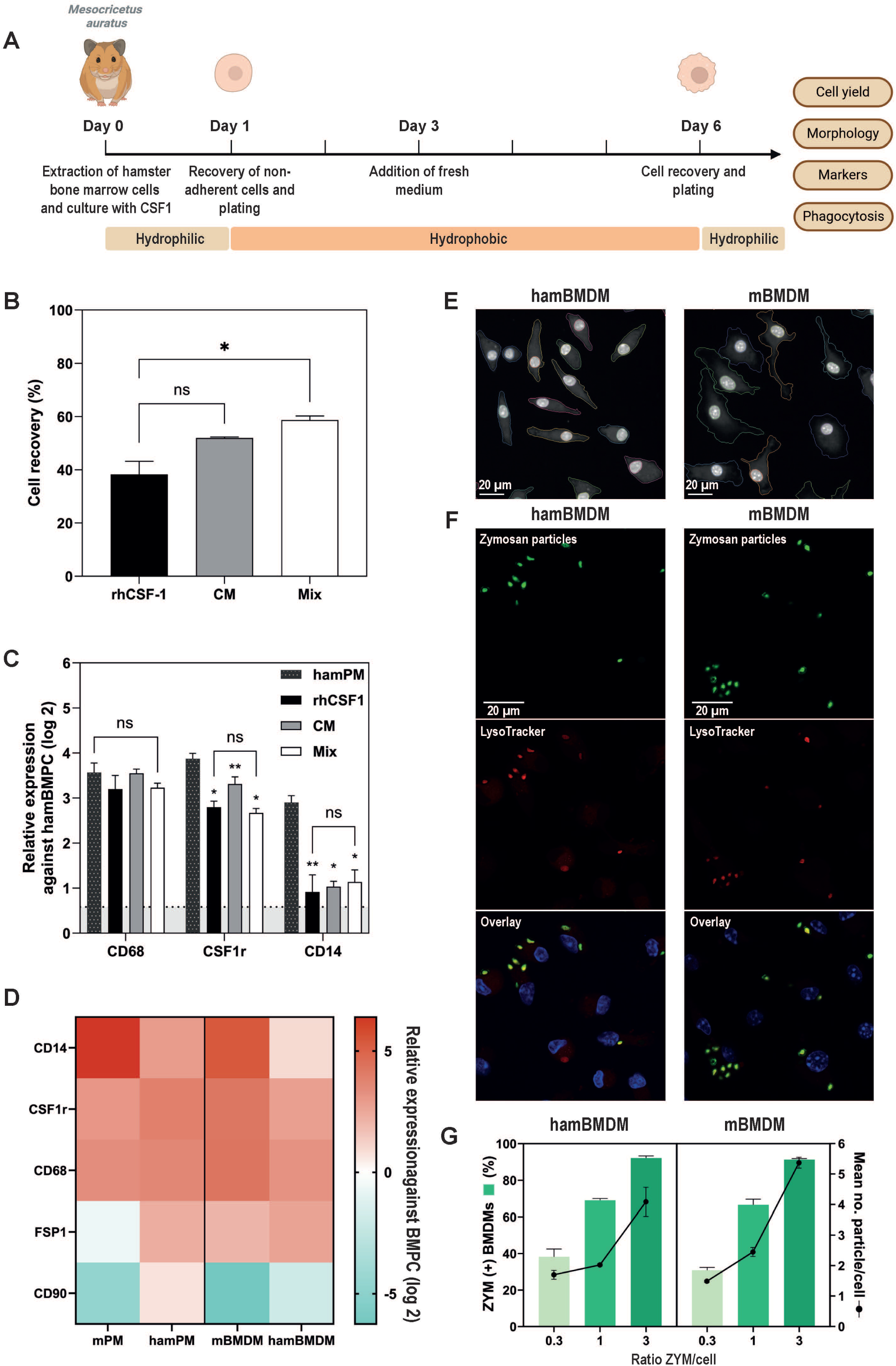
CSF1-differentiated hamster and mouse BMDMs express comparable macrophage marker profiles. (A) Overview of the experimental workflow to produce hamster bone-marrow derived macrophages. The plastic supports used for the cell culture are indicated in the orange boxes below the timeline. (B) Percent of cells recovered after 6 days of culture compared to the initial number at day 0 is shown. Results represent the mean percentage ± SEM of n = 3 independent experiments. rhCSF1, recombinant human CSF1 (black); CM, conditioned medium from baby hamster kidney cell culture (grey) Mix, combination of rhCSF1 and CM (white). Statistical analysis was made using the Tukey’s multiple comparisons test. Ns, non-significant; *p < 0.05. (C, D) Relative expression of macrophage and fibroblast markers. Total RNA was extracted from hamster or mouse bone marrow progenitor cells (ham- and mBMPC), hamster and mouse peritoneal macrophages (ham- and mPM), hamster bone marrow-derived cells after 6 days of differentiation with different sources of CSF1 (hamBMDM) and mouse bone marrow-derived macrophages (mBMDM). The expression of the indicated marker genes was monitored in 3 independent experiments by RT-qPCR. (C) Results are expressed as the mean ± SEM using hamBMPCs as calibrator. Statistical significance was evaluated using the Tukey’s multiple comparisons, p-values were calculated for each condition compared to PM. Dotted black, hamPM positive control; black, hamBMDM differentiated with recombinant human CSF1; grey, hamBMDM differentiated with BHK conditioned medium; white, hamBMDM differentiated with a mix of recombinant human CSF1 and BHK conditioned medium. Ns, non-significant; *p<0.05, **p < 0.005. (D) Heat map of relative expression levels for the indicated macrophage marker genes (CD68, CSF1r, CD14) and fibroblast marker genes (CD90 and FSP1) for the indicated cells. mPM, mouse peritoneal macrophages; hamPM, hamster peritoneal macrophages; mBMDM, mouse BMDMs; hamBMDM, hamster BMDMs. The respective mouse or hamster BMPC were used as calibrators. The color intensity indicates the mean level of relative expression (log2 fold change) as indicated by the legend. (E-G) Hamster and mouse BMDMs were collected after 6 days of differentiation and were plated overnight on glass coverslips. (E) Morphology of hamBMDMs (left image) and mBMDMs (right image). Cells were fixed with 4% PFA after overnight incubation, nuclei were stained post-fixation with Hoechst 3334 and images were acquired using an Opera Phenix Plus confocal microscope (20x magnification) and SImA script (Sequential IMage Analysis, see Methods) was applied for cytoplasm delineation for better visualization. (F) Phagocytic capacity of hamBMDMs and mBMDMs. Hamster (left panel) and mouse (right panel) BMDMs were incubated for 4 hours with Alexa-fluorescent zymosan particles. Free zymosan particles were removed by washing, phagosomes were subsequently stained using LysoTracker™ Red DND-99 and cells were fixed with 4% PFA. Images were acquired using a Leica TCS SP5 confocal microscope (63x magnification, 1.6x amplification) and were analyzed using the ImageJ software. Green, fluorescent zymosan particles (top panels); red, phagosome staining (middle panels); blue, Hoechst 3334 nuclear staining (bottom panels, overlay images are shown). (G) To assess the phagocytic capacity of hamBMDM (left panel) and mBMDM (right panel), zymosan (ZYM) particles were added to the cells at multiplicities of 0.3:1, 1:1 and 3:1 particle per cell and stained and fixed as described for panel (F). Histograms show the percentage of zymosan positive cells compared to the total number of cells counted (ZYM (+), left y-axis). Lines reflect the mean number of particles per phagocytic cell (right y-axis). Mean ± SEM from n=3 independent experiments are shown.

### Preparation of BMDM lysates for RNA extraction and real time quantitative PCR (RT-qPCR) analysis

Cells recovered after 6 days of differentiation were plated at 3×10^6^ cells per well in 6-well plates (Falcon) containing 3 ml of complete DMEM medium supplemented with 30 ng/ml of the corresponding recombinant CSF1 and incubated at 37C°, 7.5% CO2 overnight. The next day, cells were washed with warm PBS-A and scraped with 350 µl of LPB lysis buffer provided with the Macherey Nagel NucleoSpin® RNA plus kit. The recovered lysates were passed through a 1 ml syringe/26G needle and stored at -80°C until RNA extraction. The total RNA extraction was performed according to instructions provided with the Macherey Nagel NucleoSpin® RNA plus kit. Quality control of extracted RNA and reverse transcription was performed as previously described (30). RT-qPCR was performed in a final volume of 10 µl mix in an Armadillo High Performance 384-well plate PCR (Thermo Scientific) using iTaq Universal SYBR® Green Supermix (Bio-rad) and 1 µM of primers. Plates were sealed with a transparent plastic and placed in the Light Cycler® 480 system (Roche Diagnostic, Meylan, France) for 45 amplification cycles. The detailed list of primers used for the analyses is described in Table S5. Primers were designed to span introns using Ensembl transcript sequences and the LightCycler® Probe Design Software 2.0. Primer specificity was tested by NCBI blast on the corresponding rodent genome and absence of dimer formation was assessed using the melting temperature method (LightCycler ® 480 Basic Software). For each well, crossing-points were calculated based on the second derivative maximum method (LightCycler® 480 Basic Software). The mean efficacy of primers was calculated by linear regression analysis (LinRegPCR 2021.2 software) of the fluorescence raw data associated with each well (LightCycler® 480 Basic Software) (31). Wells that were not validated by LinRegPCR quality control check were excluded from further analysis. Using the NormFinder program, we selected reference genes based on their expression stability across different conditions. For macrophage and fibroblast marker analysis (Figure 1C and D, Figure S1) the reference transcripts *Gapdh/18s-RNA* (hamster samples) and *PGK1/RPS12* (mouse samples) were used. For macrophage culture adaptation (Figure 2B) the reference transcripts *actin/18s-RNA* (hamster samples) and *Mau2/RPS12* (mouse samples) were used. Relative expression was calculated using the relative expression software tool REST-MCS (32). Histograms were generated using GraphPad Prism 9.5.0. The normal distribution of the resulting log2 scale relative expression data was tested using a Shapiro-Wilk test and a Tukey’s multiple comparison test was performed for statistical analysis (p<0.05).

**Figure 2:**
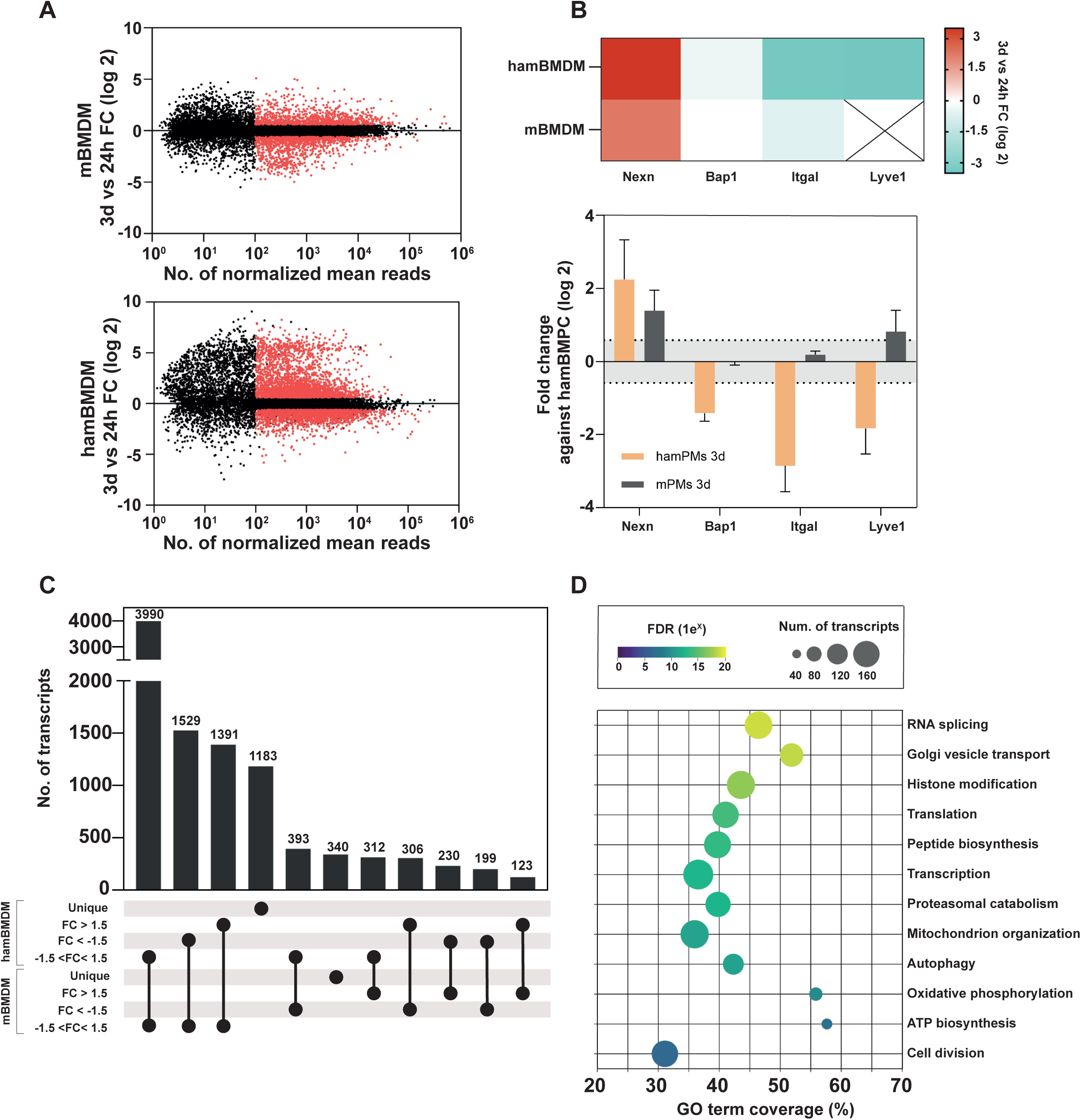
Hamster and mouse BMDMs share expression of genes implicated in core biological processes. (A) Pairwise comparison of RNA-seq analyses using three independent biological replicates of hamBMDMs (upper panel) and mBMDM (lower panel) 24 hours and 3 days after *in vitro* differentiation. Differentially regulated transcripts (absolute linear FC > 1.5, adjusted p-value (padj) < 0.05) are indicated in red. (B) Upper panel: Heatmap of differential gene expression changes during ham- and mBMDM temporal remodeling for Nexn, Bap1, Itgal and Lyve1. Lower panel: RT-qPCR validation of expression changes of the indicated genes during ham- and mPM temporal remodeling in culture. Mean values ± SEM obtained from 3 independent experiments are shown. (C) Upset plot showing the number of transcripts (histogram) ranked per group according to their expression profile. Transcripts were classified either as up-modulated (FC>1.5), down-modulated (FC<-1.5) or unmodulated (−1.5<FC<1.5) in ham- and mBMDMs. The number of transcripts (y-axis) in each group is displayed on top of their respective histogram bar. Unique transcripts were only detected in either ham- or mBMDMs. Transcripts with a normalized mean read below 100 were not considered for the analysis. (D) Functional enrichment analysis for biological processes (BP) for 3,990 unmodulated transcripts shared between hamBMDMs and mBMDMs. Each BP is plotted against its calculated GO term coverage (number of transcripts found in the datasets that share a common annotation against the background number of all transcripts with the same annotation) and represented as a circle, with the color corresponding to the false discovery rate (FDR, 1e^x^) and the bubble size to the number of transcripts in the datasets.

### Enrichment of metacyclic promastigotes by peanut agglutination

Metacyclic promastigotes were enriched as previously described (26). Briefly, promastigotes were seeded at 1.5×10^7^ parasites/ml in 15 ml of complete M199 medium at pH5.5 to induce metacyclogenesis (33). After 2 days at stationary phase parasites were collected, centrifuged for 10 min, 3000 x g at room-temperature (RT) and washed twice with PBS under the same conditions. The parasite suspension was adjusted to 1.5×10^8^ parasites/ml in incomplete M199 supplemented with 50 µg/ml of peanut agglutinin (PNA), a lectin from *Arachis hypogaea* (Sigma Aldrich). After 25 min of incubation at RT the suspension was centrifuged for 5 min, 300 x g, RT. The supernatant containing PNA-negative parasites was gently collected in a 50 ml tube filled with incomplete M199 medium and centrifuged for 10 min, 3000 x g, RT. Two subsequent washes were then performed under the same conditions, one with incomplete M199 medium supplemented with 20 mM of galactose followed by one in incomplete M199 medium only. After the last wash the pellet was resuspended in M199 medium for counting and adjustment of the parasite concentration needed for the infection.

### Phagocytosis assays with zymosan particles and *L. donovani* parasites

After 6 days of differentiation, cells were recovered and 1.5×10^5^ cells were plated on coverslips (1.5 mm) in 24 well plates (Falcon) using 500 µl of complete DMEM medium supplemented with 30 ng/ml of recombinant CSF1 (rhCSF1 or rmCSF1). For 96 well plates, 1.8×10^4^ cells were plated on PhenoPlate™ 96-well microplates using 100 µl of complete DMEM medium supplemented with 7.2 ng/ml of recombinant CSF1. Plates were incubated at 37°C, 7.5% CO2 overnight. To adapt the multiplicity of zymosan particles or parasites per cell, the number of adherent cells after the overnight incubation was estimated in control wells as follows: the culture medium of control wells was replaced by a cell lysis buffer that contains dissolved amido black 10B (0.05% w/v) in citric acid 0.1M with 1% of cetyltrimethylammonium (Cetavlon) (34) to stain the nuclei. After 5 min at 37°C, 7.5% CO2 nuclei were counted using a Malassez hematocytometer.

For phagocytosis control (Figure 1F and G), fluorescent Zymosan particles (Invitrogen™ Zymosan A *(S. cerevisiae*) BioParticles™, Alexa Fluor™ 488 conjugated) were added to the cells at increasing multiplicities of 0.3:1, 1:1, 3:1 particle per cell. Plates were centrifuged at 300 x g, 5 min, 20°C and incubated at 37°C, 7.5% CO2 for 4 hours. Coverslips were washed with warm PBS without magnesium nor calcium (PBS-A Gibco) and cells were treated with 1 µM of LysoTracker Red DND-99 (Invitrogen™) for 30 min at 37°C, 7.5% CO2 to stain the acidic vacuoles prior to fixation with 4% PFA (Electron Microscopy Sciences^TM^ - diluted from a 16% stock solution in PBS-A). Coverslips were stored at 4°C until nuclear staining and mounting on slides (see detailed protocol below).

For infection (Figure 4), *L. donovani* enriched metacyclic promastigotes or amastigotes were added to 24 or 96 well plates containing sterile glass coverslips seeded with macrophages at a multiplicity of 10:1 and 8:1 parasite per cell, respectively. Plates were centrifuged at 300 x g, 5 min, 20°C and incubated at 37°C, 7.5% CO2. After 4 hours, wells were washed with warm PBS without magnesium nor calcium (see above) to remove free parasites and placed in a new well containing 500 µl of complete DMEM medium supplemented with 30 ng/ml of their corresponding recombinant CSF1. Cells were either treated with 1 µM of LysoTracker Red DND-99 prior fixation (see above, Figure 4A) or left untreated (Figure 1B and D) and fixed with 4% PFA at 4 hours (D0), 24 hours (D1), 48 hours (D2), 72 hours (D3) and 144 hours (D6) to be stored at 4°C until nuclear staining, mounting on slides and image acquisition (see below). For long-term infection, cells on coverslips were infected with 10:1 *mCherry* positive *L. donovani* amastigote parasites and fixed after days 7 and 21 before image acquisition using the EVOS™ M5000 Imaging System (40x).

For hamBMDM and mBMDM infection and to follow parasite burden, all experiments were performed in 96-well plates applying automated image acquisition using the confocal microscope Opera Phenix Plus confocal microscope (20x magnification). Fifteen fields per well were captured with a 5% overlap to avoid field-edge artifacts. This setup enabled analysis of over 1,500 cells per well. For three biological replicates and five technical replicates per condition, more than 7,500 cells were analyzed per condition. Parasites were then automatically counted to estimate the burden (see section ‘Automatic counting of parasites using SImA).

### BMDM and parasite staining

For phagocytosis of zymosan particles (Figure 1F and G) and infection by *L. donovani* (Figure 4), an immunostaining of parasites on fixed cells was performed prior to the Hoechst 33342 nuclear staining. Coverslips were washed in their wells with 1 ml of PBS and incubated 5 min at room temperature. PBS-A was removed, and aldehyde groups were blocked for 20 min at room temperature using 1 ml of NH4Cl 50 mM. Non-specific antibody interactions were prevented with PBS, saponin 0.1 mg/ml supplemented with 10% goat serum for 15 min at room temperature (RT). Coverslips were then incubated at RT for 45 min in PBS supplemented with 2.5% gelatin and a 1/1000 dilution of immune serum from *L. donovani*-infected hamsters, washed 3 times successively with PBS before a 45 min incubation at RT with 5 µg/ml of a goat anti-hamster IgG conjugated to Alexa Fluor 488 (Jackson ImmunoResearch) in PBS with 2.5%. Coverslips were washed three times and cell nuclei (BMDMs and parasites) were stained in PBS supplemented with 0.1% Saponine (Sigma Life Science) and Hoechst 33342 at a final concentration of 5 µg/ml (ThermoFisher Scientific) during 10 min at RT. Lastly, coverslips were washed with 1 ml of PBS with 0.1% Saponine (Sigma Life Science) for 5 min at RT, rinsed in distilled water (Gibco), mounted on slides with Antifade Mounting Medium (Vectashield® - Vibrance™) and incubated at room temperature overnight. After 24 hours, images were acquired with the Leica TCS SP5 confocal microscope.

For parasite infection assays in 96-well plates, we adapted the previously described protocol for immunostaining on plate. After the last washes, 200 µl of PBS was added to each well and plates were stored at 4°C until image acquisition.

### Automatic counting of parasites using Sequential Image Analysis (SImA)

In collaboration with the BioImagerie Photonique (UTechS PBI) platform at Institut Pasteur we developed a script for automated quantification of parasites within hamster and mouse macrophages using SImA, a python software package designed for quantitative analysis of images taken from fluorescence microscopy (Table S6). Host cell nuclei were detected by Hoechst 33342 staining (Figure S5, image 2), and cytoplasmic boundaries were defined using the background noise from the Alexa 488 staining (Figure S5, image 3). Hamster and mouse cells located at the borders of acquired fields were excluded from downstream analysis to avoid bias in parasite burden counting and script complications (Figure S5, image 4). Nuclear areas ≤ 200 µm² were classified as described by Swanson et al. (35), while those > 200 µm² were considered contaminating cells (e.f. fibroblasts) (36). Hoechst staining was used to identify the kinetoplasts of parasites located within the macrophage parasitophorous vacuole, excluding the nuclear region of the macrophage (Figure S5, panel 5). Based on this analysis, cells were categorized as either uninfected or infected (Figure S5, panel 6). Infected cells were further classified according to the degree of intracellular parasite burden, allowing us to track the distribution of intracellular parasites over time in different macrophage classes (Figure S5, panel 7).

### BMDM cultures for RNA-seq analysis

After 6 days of differentiation hamster and mouse BMDMs were recovered and 2×10^6^ cells were plated in 12 well plates (Falcon) using 2 ml of complete DMEM medium supplemented with 40 ng/ml of recombinant CSF1 (rhCSF1 or rmCSF1). Plates were incubated at 37°C, 7.5% CO2 overnight. The next day, cell medium was replaced with 2 ml of fresh, pre-warmed, complete DMEM medium supplemented with 40 ng/ml of recombinant CSF1. Cells were lysed after 24 or 72 hours, and the RNA was extracted as previously described (see above ‘Preparation of BMDM lysates for RNA extraction and real time quantitative PCR (RT-qPCR) analysis’).

### RNA quality control, RNA sequencing and transcriptomic analysis

RNA quality control was performed as previously described (37). Briefly, the quality of RNA extracts was assessed by optical density measurement using Nanodrop (Kisker, http://www.kisker-biotech.com) and by electrophoresis on a Lab-on-a-chip product using the Agilent 2100 Bioanalyzer (Agilent, http://www.chem.agilent.com). The RNA integrity number (RIN) of every sample was above 9 out of 10.

Libraries were prepared from 250 ng of RNA using Illumina Stranded mRNA Prep (Illumina) following the manufacturer’s protocol. Briefly, mRNA was captured by oligo-dT magnetic beads, fragmented and reverse transcribed. Sample-specific barcodes were ligated to the cDNA which was then amplified by 15 cycles of PCR. Note only the first cDNA strand was amplified. Purification of unbound adaptors and primers was done on AMP beads (Beckman Coulter) added to samples in a 0.8:1 ratio. The resulting stranded libraries comprised fragments from 200 to 1000 bp with a peak around 490 bp as visualized on a 5300 Fragment Analyzer (Agilent Technologies). Libraries were pooled, purified on AMP beads as above, diluted to 0.8 nM and sequenced on a NextSeq 2000 system (Illumina) using a P3 50-cycle sequencing kit. The yield was 1200 mln 67-bp single-end reads (30 – 60 mln reads per sample).

RNA-seq analysis was performed with Sequana v0.15.4 (38). Specifically, we used the RNA-seq pipeline v0.18.1 (https://github.com/sequana/sequana_rnaseq), which is built on top of Snakemake 7.3.2 (39). The pipeline was executed with default parameters and reproducible containers from the Damona project (https://github.com/cokelaer/damona). Reads were mapped to the golden hamster (*Mesocricetus auratus*, MesAur1.0-v100) or to the mouse (*Mus musculus*, GRCm39 v105) genome assembly from Ensembl using STAR 2.7.3a (40). A dual-reference analysis was performed by concatenating the *Leishmania donovani* genome. Gene-level quantification was conducted using FeatureCounts 2.0.0 (41) assigning reads to genomic features based on the MesAur1.0-v100 or the GRCm39 annotation only, while accounting for strand-specificity information. Differential expression analysis was performed using DESeq2 v1.24.0 (42) scripts available in Sequana library. Statistical testing identified differentially expressed genes by comparing samples, with significance determined using Benjamini-Hochberg adjusted p-values (false discovery rate FDR < 0.05). For gene enrichment, we established orthologous relationships between *Mesocricetus auratus* and *Mus musculus* by extracting Ensembl gene IDs from the annotation. Ortholog mapping to *Mus musculus* was performed using g-profiler (43).

### Gene Ontology Analysis

We performed gene ontology (GO)-enrichment analyses on transcript datasets using the STRING plugin of the Cytoscape software package (v3.10.2) under Java 17.0.5. Transcripts with normalized mean read below 100 were removed from the study. A transcript was considered differentially expressed when its linear fold change was above 1.5 and its adjusted p-value (padj) below 0.05. STRING networks were constructed for each dataset using the *Mus musculus* as reference and a confidence score cutoff of 0.4. Functional enrichment results were visualized, showing the corresponding false discovery rates (FDR) and the absolute number of transcripts annotated for each biological process (Figure 2D). We manually curated the list of biological processes to avoid redundancy among closely related terms and to exclude overly broad categories. The GO term coverage was calculated as the number of transcripts identified in our dataset relative to the background number of transcripts annotated in Cytoscape. Enrichment maps were generated using g:Profiler output with a node cutoff 0.05, edge cutoff 0.5 and data set edges – automatic sparse 1/5. Terms and bubble were manually curated and positioned for clarity (Figure S3).

### Polarization marker list

A list of M1 and M2 markers (Table S4) established in Zhang et al. 2024 (44) and based on results from Orecchioni et al. 2019 (45) and Jablonski et al., 2015 (46) was used.

### Software for graphic representation

The website ChiPlot (https://www.chiplot.online/) was used to plot Figure 2D, Figure 3B and D and Figure 4B and D. The software GraphPad Prism 10.2.3 was used for Figure 1B and D and G, Figure 2A and C, Figure 3A, Figure 4C and D and Figure S1, Figure S3B and C. Figure 1A and Figure S3A were created with BioRender.com. Figure S2 was generated using Cytoscape software package (v3.10.2).

**Figure 3:**
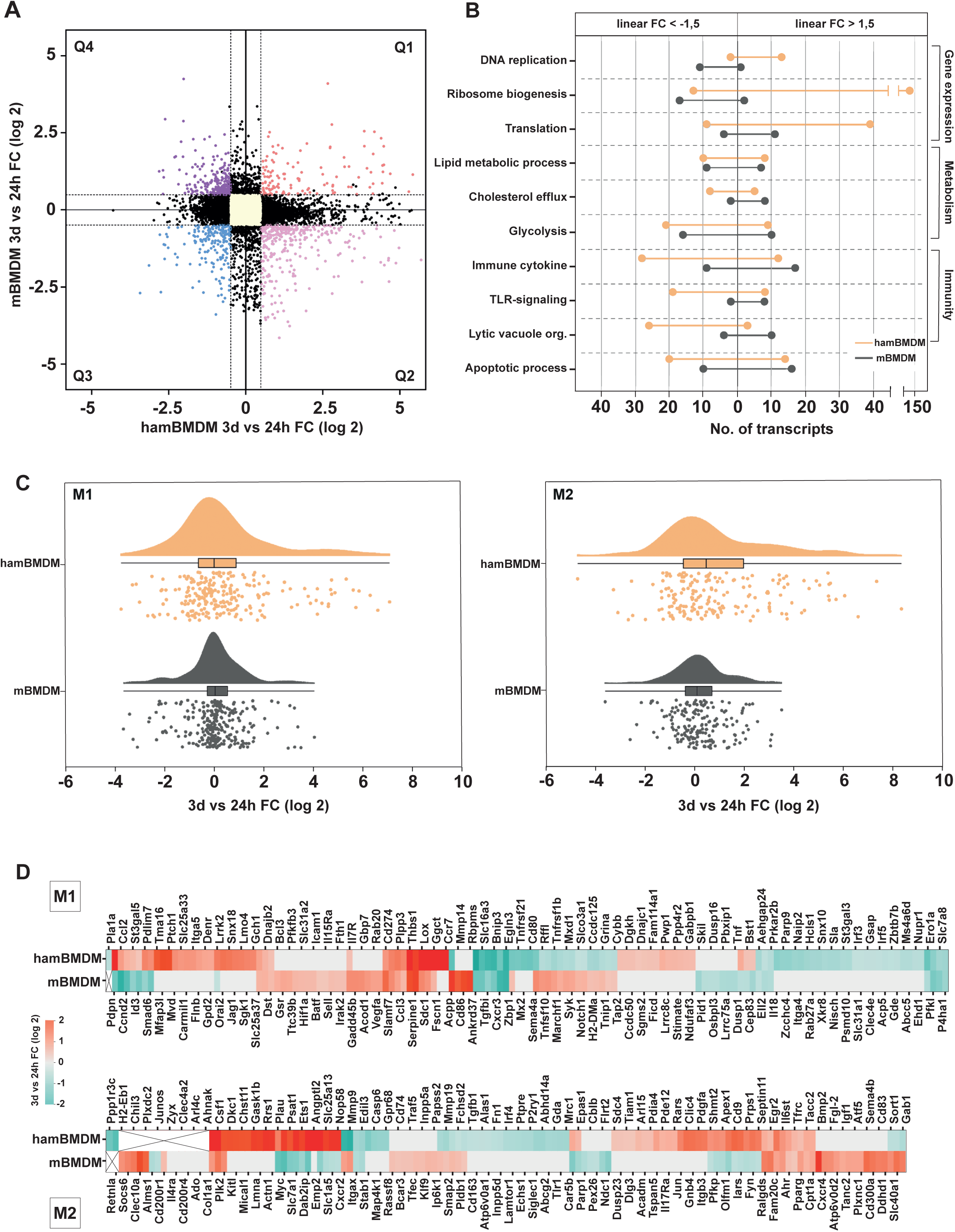
Hamster and mouse BMDMs show rodent-specific expression changes during *in vitro* temporal remodeling. (A) Double ratio plot showing the log2 FC of each transcript (dots) between expression changes observed in hamBMDMs (x-axis) and in mBMDMs (y-axis) at 3 days vs 24 hours. Transcripts are considered differentially regulated when absolute linear FC > 1.5 and adjusted p-value (padj) < 0.05. Dotted lines (in black) correspond to the absolute linear FC > 1.5 threshold. Transcripts with a normalized mean read below 100 were not considered for the analysis. According to expression changes observe in the rodent systems, transcripts clustered into four quadrans (Q) and were colored according to their expression trends. Yellow, unmodulated transcripts; red, transcript showing increased abundance during culture of both ham- and mBMDMs (Q1); blue, transcript showing decreased abundance during culture of both ham- and mBMDMs (Q3); pink, transcript showing increased abundance in hamBMDM and decreased abundance in mBMDMs (Q2); purple, transcript showing decreased abundance in hamBMDM and increased abundance in mBMDMs (Q4). (B) Dumbbell representation showing the GO-terms (y-axis) and the corresponding number of transcripts (x-axis) that are up (right side) or down regulated (left side) during hamBMDM (orange) and mBMDM (grey) temporal remodeling. (C) Raincloud plots showing the distribution of hamBMDMs and mBMDMs polarization markers (see Table S4). Transcripts were sorted into M1 (left panel) or M2 (right panel) polarization markers and plotted based on their FC values (x-axis) for hamBMDMs (orange) and mBMDMs (grey). Each dot corresponds to a transcript as a function of its log2 FC (rain, lower part). Histograms (cloud, upper part) represent the distribution of the transcript FC in the datasets based on kernel density estimate. Boxplots (middle part) indicate the 1^st^ quartile, the median and the 3^rd^ quartile of the datasets. (D) Heatmap of individual expression changes of M1 (upper panel) and M2 (lower panel) markers during temporal remodeling of hamBMDM (upper row) and mBMDMs (lower row).

## Data availability

Datasets are available on the EMBL-EBI website in the ArrayExpress-ENA data collection under the following accession number: E-MTAB-15420

## RESULTS

### Hamster bone marrow-derived, macrophage-like cells (hamBMDMs) express macrophage characteristics and show high phagocytic capacity

The large-scale production of quiescent macrophages is essential for many systems-level analyses that require abundant input material. Instead of isolating primary macrophages from tissues, which can alter their activation state, a more reliable and reproducible approach involves differentiating bone marrow-derived macrophages (BMDMs) from bone marrow progenitor cells (BMPCs) using colony-stimulating factor 1 (CSF1). While well-established protocols exist for generating murine (m)BMDMs, adapting this method to other rodent models is often hindered by the lack of commercially available autologous CSF1. This challenge is particularly evident in hamsters, a key animal model for studying immune responses and infectious diseases affecting humans, such as visceral leishmaniasis. Here, we adapted the mBMDM protocol (29) for hamster (ham)BMPCs and evaluated alternative CSF1 sources for their ability to support hamster (ham)BMDM differentiation (Figure 1A).

Culture of hamBMPCs in presence of *E. coli*-produced recombinant mouse (rm)CSF1 or recombinant human (rh)CSF1 resulted in low cell recovery of hamBMDMs compared to mBMDM controls (data not shown). In contrast, robust hamBMDMs recovery was observed after 6 days of differentiation using 100 ng/ml rhCSF1 produced in the eukaryotic human embryonic kidney cell line (HEK 293), indicating that post-translational modifications might increase the activity of heterologous CSF1 (Figure 1B). Similar levels of cell recovery were attained with conditioned medium (CM) of baby hamster kidney (BHK) cells, which was further increased to 60% using a mixture of 100 ng/ml rhCSF1 and 5% CM.

The differentiation state of hamBMDMs treated with rhCSF1, CM or a mix of both was first assessed by RT-qPCR determining the relative expression of the macrophage markers CD68, CSF1 receptor (CSF1r) and CD14 (47–49) using hamBMPCs as calibrator. Mature tissue-resident macrophages recovered from the hamster peritoneal cavity (hamPMs) were used as positive controls (Figure 1C). Irrespective of the differentiation condition, CD68 and CSF1r expression was induced in hamBMDMs to levels similar to hamPMs. In contrast, the level of CD14 expression was more than twice as high in the hamPM control population compared to hamBMDMs. Even though CD14 is described as a monocytic marker, it has been reported to be heterogeneously expressed among macrophage subpopulations, with PMs expressing a higher level compared to for example alveolar macrophages (50).

To provide a clearer contrast for the expression of macrophage markers in hamBMDM culture, we used BHK as a non-hematopoietic calibrator, which allowed the comparison of hamBMPCs with hamBMDMs. We observed that hamBMPCs were already expressing a high level of CD14 (8-fold increase, Figure S1A), explaining the mild enrichment observed for this marker compare to CD68 and CSF1r (Figure 1C). For subsequent analyses, we selected the rhCSF1-based differentiation protocol, as it closely aligns with the mBMDM protocol and avoids potential side effects of BHK-conditioned medium, such as the presence of additional hematopoietic growth factors (51) that may influence macrophage markers, cytokine production, and metabolism (52, 53).

To further validate our systems, we next compared the expression profile of our hamBMDMs and hamPMs with mBMDMs and mPMs for the macrophage and fibroblast markers shown in Figure 1C to assess potential cell contamination that may affect the macrophage differentiation status during culture (54, 55) (Figure 1D). In comparison to their respective PMs, mBMDMs and hamBMDMs presented a similar profile of expression for the macrophage markers CD68, CSF1r and CD14. Likewise, the profiles of fibroblast markers FSP1 and CD90 showed equal or reduced signals in mBMDMs and hamBMDMs compared to the mPMs and hamPMs controls, indicating only minor fibroblast contamination (Figure S1D, Figure 1B). Both hamBMDMs and mBMDMs showed macrophage-like morphology, characterized by a flat and irregularly shaped body with smooth edges and a round-to oval-shaped nucleus (Figure 1E), features that they share with human macrophages differentiated from peripheral blood mononuclear cells (PBMCs) (56, 57) and hamster macrophages isolated from the spleen (58).

We next evaluated hamBMDM and mBMDM phagocytic capacity as one of the hallmarks of macrophage biology using fluorescent zymosan particles. Zymosan was efficiently ingested by hamBMDMs into lysotracker-positive vacuoles at similar levels to those of mBMDM as monitored by microscopy (Figure 1F). Quantitative analysis of phagocytic capacity using increasing particle-to-cell ratios (0.3, 1, 3) confirmed these results, with at least one particle observed in 92% of both hamBMDMs and mBMDMs at the lowest zymosan concentration (Figure 1G).

Altogether these results validate hamBMDMs differentiated in presence of rhCSF1 as *bona fide* macrophages that reproduce morphology, gene expression and phagocytic capacity of mBMDMs.

### Comparative transcriptomic profiling of hamster and mouse BMDMs reveal a common core transcriptome but rodent-specific gene expression changes during *in vitro* maturation

We next applied RNAseq on ham- and mBMDMs collected at 24 hours and at 3 days post-differentiation to evaluate temporal changes (referred to as ‘remodeling’ or ‘maturation’) in gene expression within each BMDM system (3 days vs 24 hours) and changes between the BMDM systems (hamster vs mouse).

In mBMDMs, 15,733 transcripts were detected (x-axis) of which 2,146 showed changes in abundance during maturation (Figure 3A, highlighted in red, abs. FC>1.5, adj p-value<0.05, normalized mean reads>100). Among these, 958 (45%) were up-regulated and 1,188 (55%) were down-regulated. Gene-ontology (GO) enrichment analysis of up-regulated transcripts reveals enrichment in the GO terms ‘inflammatory response’ and ‘cytokine production’ in mature mBMDMs, while down-regulated transcripts were enriched for the GO terms ‘DNA repair’ and ‘cell cycle transition’ (Table S1).

Likewise, the analysis of hamBMDMs detected 14,250 transcripts of which 5,186 were found differentially modulated during maturation, with 2,701 (52%) showing increased and 2,485 (48%) decreased transcript abundance (Figure 2A). Enrichment analysis identified the GO terms ‘RNA processing’ and ‘protein production’ for the up-regulated transcripts, and the GO terms ‘vacuole organization’ and ‘defense response related processes’ for the down-regulated transcripts, among others (Table S1).

However, unlike mBMDMs, mature hamBMDMs showed strong increased expression (log2 FC>5) for 300 transcripts (Figure 2A), which were enriched in the GO terms ‘extracellular matrix (ECM) organization’ implicating collagen fibril as well as angiogenic factors (Table S1), suggesting a propensity of hamBMDMs for tissue-mending functions characteristic of M2 polarized macrophages.

To test if these differences between hamBMDMs and mBMDMs are independent of our differentiation protocol, we examined whether these expression changes were recapitulated in culture using terminally differentiated peritoneal macrophages (PMs) isolated from the peritoneal cavities of mice and hamsters. We selected 4 genes (Nexn, Bap1, Itgal and Lyve) from our transcriptomic dataset based on their distinct expression profiles between 24 hours and 3 days in ham- and mBMDMs (Figure 2B, upper panel) and assessed by RT-qPCR their expression levels over time in hamPM and mPM populations. Strikingly, the regulation of the selected genes in the two distinct rodent PMs closely mirrored the expression patterns observed in their BMDMs counterparts (Figure 2B, lower panel), revealing that the differential expression program identified in our transcriptomic analyses can be, at least partially, extended to resident macrophages and thus represent intrinsic, species-specific macrophage signatures.

Direct comparison of the ham- and mBMDM expression changes revealed 3,990 transcripts that remained unmodulated over time in both macrophage systems (Figure 2C). This core of constitutive transcripts is enriched in GO terms for basic biological processes such as ‘transcription’, ‘translation’ or ‘cell division’ (Figure 2D). However, significant expression differences were identified for almost 49% of the transcripts (fold change > 1.5 and adj. p-value < 0.05), including 340 mouse-specific and 1,183 hamster-specific transcripts (Figure 2C), suggesting intrinsic and rodent-specific differences during maturation, which are investigated in detail below.

### Hamster BMDMs show characteristics of M2 polarized macrophages

To further investigate interspecies differences, we plotted the ratios of expression changes (day 3 versus 24 hours post-infection) observed in our ham- and mBMDMs transcriptomics datasets (Figure 3A, Table S2). Central transcripts (in yellow) correspond to core unmodulated transcripts previously analyzed (see Figure 2D). Each quadrant delineates a population of transcripts that evolved either the same way (Q1 and Q3) or inversely (Q2 and Q4) in the ham- and mBMDMs over time. We first searched for any functional enrichment within each quadrant (Figure S2A-B, Table S3). Transcripts that were up-modulated in both mature mBMDMs and hamBMDMs (Q1) are enriched in GO terms such as ‘proteasomal protein catabolic process’ and ‘regulation of translation’. In contrast, those that were down-modulated in both systems (Q3) show enrichment in GO terms ‘DNA repair’ and ‘ATP metabolic processes’. Transcripts up-modulated in mBMDMs but down-modulated in hamBMDMs (Q4) are associated with the GO terms ‘autophagy’, ‘vacuole organization’ and ‘regulation of NFκB signaling’. Conversely, transcripts down-modulated in mBMDMs but up-modulated in hamBMDMs (Q2) are enriched in ‘mRNA processing’, ‘ribosome’ and ‘peptide biogenesis’.

We next mined our data sets for signals that are distinct between the two rodent systems and may be linked to their different degree of susceptibility to VL. A number of key biological processes implicated in gene expression, metabolism and immunity were identified (Figure 3B). Overall, the total number of transcripts modulated across these processes is increased in hamBMDMs compared to mBMDMs (Figure 3B), with the most strongly upregulated transcripts showing enrichment for the GO terms such as ‘DNA replication, ‘translation’ and ‘ribosome biogenesis’. Thus, hamBMDM may be biosynthetically more active compared to mBMDM, potentially providing a nutritionally richer niche to support intracellular *Leishmania* proliferation.

In contrast to mBMDMs, hamBMDMs show reduced expression of transcripts annotated for the GO term ‘cholesterol efflux’, suggesting intracellular accumulation of cholesterol, an essential membrane component known to support parasite growth (59–61). Likewise, hamBMDMs showed reduced expression for transcripts annotated for GO terms linked to innate, anti-microbial resistance and induction of the inflammatory response, such as ‘lytic vacuole organization’ or ‘TLR signaling’ (Figure 3B), further supporting the notion that hamBMDMs may be more permissive for *Leishmania* infection. This is further supported by investigating expression signatures characteristic for the different macrophage polarization states. Based on a curated list of polarization markers (Zhang, 2024), mature hamBMDMs and mBMDMs showed similar expression patterns for M1 marker genes (Figure 3C). In contrast, a clear bias towards M2 polarization was observed in hamBMDMs as indicated by the significant increase in the median expression change across all M2 marker genes (0.37 versus 0.11 log2-fold change in respectively hamBMDMs and mBMDMs). Direct comparison of M1 and M2 marker gene expression between the rodent systems further confirms this bias (Figure 3D) and revealed that the expression changes observed for both types of polarization markers showed only little overlap in ham- and mBMDMs (13% and 10% for M1 and M2 maker genes, respectively).

In conclusion, our results suggest that mature hamBMDMs express more favorable conditions for intracellular *Leishmania* growth compared to mBMDMs, a possibility we investigated in the following using our *in vitro* macrophage infection assay.

### Hamster BMDMs are more permissive to *L. donovani* parasites compared to mouse BMDMs

We first set up an experimental system to quantify the capacity of hamBMDMs to phagocytose *L. donovani* parasites. Infectious, metacyclic promastigotes were enriched by agglutination from day 2 stationary culture and added at a ratio of 10:1 to day 6 differentiated ham- and mBMDMs. Free parasites were removed after 4h, and parasite uptake and localization to the PV was visualized after 24h of infection by fluorescence microscopy using a parasite-specific immune serum and the lysosomal fluid phase marker LysoTracker Red DND-99 (Figure 4A). We next applied this protocol using (i) *L. donovani* amastigotes purified from the liver of infected hamsters, (ii) amastigote-derived, virulent promastigotes from early culture passage (p2) at day 2 of stationary growth phase that were either enriched or not for metacyclics, and (iii) attenuated promastigotes from late culture passage (p20) at day 2 of stationary growth phase (62) (Fig. S3A). While all parasites tested were internalized by both ham- and mBMDMs, a statistically significant increase in uptake was observed in hamBMDMs for p2 metacyclic, p2 stationary-phase and p20 stationary-phase parasites (Figure 4B), further supporting the notion that hamBMDMs may be more suitable host cells for *Leishmania* infection.

**Figure 4:**
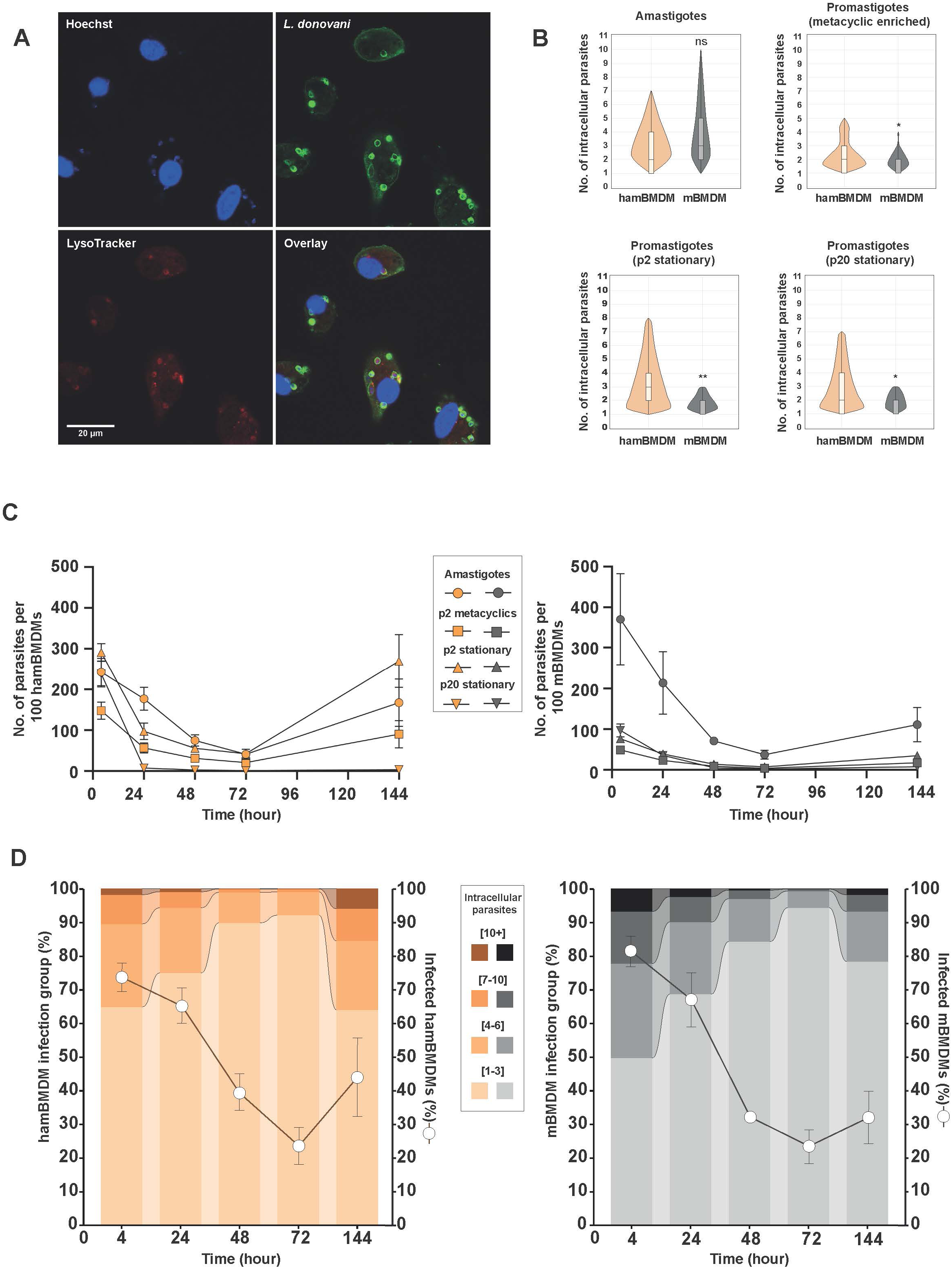
hamBMDMs are more permissive to *L. donovani* intracellular proliferation compared to mBMDMs. Macrophages were incubated for 4 hours with *L. donovani* promastigotes at an MOI of 10 parasites per macrophage (10:1) (p2 metacyclic, p2 stationary and p20 stationary), or amastigotes isolated from hamster spleen at an MOI 8:1. After 4, 24, 48, 72 and 144 hours cells were fixed with 4% PFA and nuclei of cells and parasites were stained using Hoechst 33342. Images were acquired using an Opera Phenix Plus confocal microscope (20x magnification). Intracellular parasites were automatically counted applying SImA (see Methods) on parasite cell body and nuclear staining. (A) At 24h post-infection with *L. donovani* metacyclic-enriched promastigotes, hamBMDMs were stained with LysoTracker™ Red DND-99 to validate the presence of intracellular parasites within the phagolysosome. Parasites were stained using hamster immune serum. Images were acquired using a Leica TCS SP5 confocal microscope (63x magnification, 1.6x amplification) and analyzed using the ImageJ 1.54g software under Java 1.8.0_345. (B) Phagocytosis capacity of hamBMDMs (orange) or mBMDMs (grey) incubated with *L. donovani* promastigotes and amastigotes. The repartition of intracellular parasites was determined automatically applying SImA on BMDM and parasite nuclear staining. Results are representative of n=3 independent experiments. (C) Evaluation of ham- and mBMDM parasite burden after infection with promastigotes and amastigotes according to the legend shown in the graph. The number of intracellular parasites for 100 hamBMDMs (left panel) and mBMDMs (right panel) (y-axis) is reported over time (x-axis). (D) Proportion of intracellular parasites per infected hamBMDMs (left) or mBMDMs (right). For all time-points of infection (x-axis), infected hamBMDMs (left panel) and mBMDMs (right panel) were grouped according to intracellular parasite burden (left y-axis) as indicated in the legend. The percentage of infected BMDMs against the total number of BMDMs counted is projected as a line plot (right y-axis).

Following intracellular parasite burden until day 6 post-infection confirmed this result. While both ham- and mBMDMs eliminated a fraction of parasites during the first 72 hours post-infection, intracellular parasite burden showed a more robust recovery thereafter in hamBMDMs for amastigote and p2 parasite infections. No recovery is observed for the attenuated p20 in either rodent system, revealing that hamBMDMs maintain an anti-microbial potential comparable to mBMDMs.

While the percentage of infected macrophages over time is similar in ham- and mBMDMs (Figure S3B), their respective intracellular parasite burden shows important differences (Figure 4D). We deconvoluted the parasite burden observed for the infection with amastigotes by defining four infection groups corresponding to low (1-3 parasites per infected cell), medium-low (4-6 parasites), medium-high (7-10 parasites) and high (>10 parasites). Even though more mBMDMs were infected at higher parasite burden at 4 hours post-infection compared to hamBMDMs, only the latter one showed a robust increase in intracellular parasite growth with 36 % of the cells harboring 4 or more parasites, compared to only 22% the in mBMDMs. Of note, the percentage of infected macrophages increased in both ham- and mBMDMs, which is not caused by re-infection of extracellularly growing parasites (whose absence was monitored microscopically during the infection period) but due to the loss of uninfected cells and thus enrichment of only infected ones.

*Leishmania* parasites indeed are known to prolong the lifespan of host macrophages, exploiting them as niches for sustained proliferation (63). This is confirmed in our system: after three weeks of infection, hamBMDMs were viable and exhibited a heavy parasite burden, showing balloon-shaped expansions laden with amastigotes. In contrast, mBMDMs effectively restricted parasite proliferation, maintaining normal cellular morphology and lower intracellular parasite load. These preliminary *in vitro* observations recapitulate the *in vivo* infection outcomes, where hamsters develop progressive VL while mice control parasite growth more effectively.

Altogether these results validate the potent microbicidal properties of hamBMDMs as they reduced parasite burden of virulent amastigotes and p2 promastigotes during the first 3 days of infection and completely cleared attenuated p20 parasites. At the same time, compared to mBMDMs they are more permissive to intracellular infection with virulent *L. donovani*, establishing a link between their M2-like immuno-metabolomic phenotype and their increased susceptibility, which may contribute to symptomatic VL observed in hamsters.

## Discussion

A central unresolved question in *L. donovani* infection is why some individuals are resistant while others develop severe disease despite no apparent immunological, metabolic, or nutritional deficiencies. We addressed this important open question by comparing *L. donovani* susceptible, hamster bone marrow-derived macrophages (hamBMDMs) with resistant mouse (m)BMDMs. Optimizing and validating an experimental protocol for generating bulk quantities of hamBMDMs allowed us to reveal intrinsic and rodent-specific differences in macrophage biology by genome-wide transcript profiling that correlated with increased parasite burden observed in hamBMDMs and may influence the opposite outcomes of visceral leishmaniasis in these rodent systems.

During *in vitro* infection, the first physical interaction between *Leishmania* and macrophages is ligand-receptor dependent triggering host signaling pathways and immune responses. Many host cell receptors have been described to interact with either promastigote or amastigote surface ligands or bound opsonins, such as Fc gamma receptors (FcγRs), fibronectin receptors (FnRs), complement receptors (CRs), toll-like receptors (TLRs) or the mannose receptor (MR) (64). Three of these receptors - FcγRs, TLR9 and MR - showed reduced expression in mature hamBMDMs compared to mBMDMs. While MR does not appear to play a critical role in the response to *Leishmania* infection (Akilov, 2007), and FcγR engagement by IgG-coated *Leishmania* often modulates cytokine expression in macrophages toward an anti-inflammatory profile that favors parasite survival (64, 65), TLR9 plays a well-established role in anti-leishmanial immunity. It promotes dendritic cell activation, proinflammatory cytokine production and a protective Th1 response (66, 67). Therefore, the downregulation of TLR9 expression in hamBMDMs may be a first intrinsic aspect that contributes to enhanced *L. donovani* survival.

Our data reveal polarization as a second intrinsic difference between ham- and mBMDMs. A hallmark of macrophages is their phenotypic plasticity to adopt specialized functions through polarization into pro-inflammatory and anti-microbial M1, or anti-inflammatory and tissue-repairing M2 cells (45, 68). While this specialization is usually governed by specific cytokines, our study revealed the intrinsic capacity of both ham- and mBMDMs to mature during three days of culture towards a mixed polarization profile when compared to immature cells at 24 hours after differentiation. In particular, we show that hamBMDMs are skewed toward an M2-like polarization state that supports intracellular *Leishmania* survival. Mature hamBMDMs showed increased expression of the M2 marker genes Arg1 and ODC1, which direct the L-arginine catabolism toward polyamine biosynthesis instead of NO production (69). In addition, mature hamBMDMs show increased expression of the arginine transporters SLC7A1 and SLC7A6, which are respectively down-modulated or unchanged in mBMDMs. This expression profile likely increases the pool of L-arginine, which then will be available for salvage by *Leishmania* that is auxotroph for this essential amino acid, thereby enhancing intracellular parasite proliferation (70).

Finally, the gene expression profile of hamBMDMs reveals downregulation of genes associated with cholesterol efflux during hamBMDM maturation that are not affected in mBMDMs. Species-specific differences are common in cholesterol homeostasis and have for example been observed between human and murine cells for the expression of the VLDL receptor that mediates lipoprotein uptake (71), or Liver X receptors (LXRs) that can promote cholesterol efflux (72, 73). Compared to mouse macrophages, mature hamster macrophages show lower NR1H2 (LXRβ) expression, while mouse macrophages have reduced expression of the master transcriptional regulator of the cholesterol pathway SREBF2. This may lead to cholesterol accumulation in hamster macrophages that may benefit *L. donovani* survival. Indeed, similar to their auxotrophy for L-arginine, *Leishmania* parasites are unable to synthesize cholesterol de novo (74) and must scavenge it from the host cell to support membrane biogenesis essential for their growth. The critical role of host cell cholesterol in facilitating intracellular *Leishmania* infection has been well established for *L. amazonensis* (75–78). Likewise, *L. donovani* is a cholesterol auxotroph and needs to acquire this essential lipid from the host cell, for example by depleting it from the macrophage membrane (79). Increased host lipid uptake indeed correlates to enhanced intracellular proliferation as observed in a drug resistant *L. donovani* isolate from India (80). As a key membrane component, cholesterol may also explain the reduced uptake of promastigotes by mouse macrophages we observed in our study, since its depletion has been shown to impair promastigote but not amastigote entry (81).

In conclusion, our study highlights intrinsic, species-specific differences in macrophage biology that may potentially contribute to the contrasting susceptibility of hamster and mouse macrophages to *L. donovani* infection. The parasite’s success in establishing infection depends on its ability to adapt to the intracellular environment and subvert host processes that normally restrict pathogen survival within the phagolysosome (6, 82, 83). Our findings indicate that hamBMDM may be particularly vulnerable to these subversion strategies as they already provide inherent biological characteristics to support *L. donovani* infection. These traits together with the absence of inducible nitric oxide synthase (iNOS) expression due to a defective promoter region (84, 85) may be sufficient to explain the hamster’s susceptibility to develop lethal VL. Given the physiological and metabolic similarities between hamsters and humans, particularly in lipid metabolism and infectious disease pathophysiology, hamBMDMs represent a powerful cellular system to identify infection-associated signals *in vitro,* which can then be assessed *in vivo* by experimental hamster infection. Our hamBMDM protocol thus offers a valuable approach for generating systems-level insights relevant to human *Leishmania* infection. Moreover, it can serve a blueprint for studying other intracellular pathogens of clinical relevance, such as *Mycobacterium* spp. and SARS-CoV-2, for which hamsters serve as a model for human infection. The combined use of murine and hamster macrophage systems provides a powerful platform to dissect the molecular mechanisms underlying pathogen survival and host resistance, paving the way for comparative transcriptomic, metabolomic, and loss-of-function studies that can uncover how *L. donovani* and other microbes exploit host-specific macrophage programs to establish chronic infection.

## Acknowledgments

This work was supported by Université Paris Cité through a three-year PhD fellowship within the BioSPC Doctoral School (ED 562). Additional support was provided by the ANR Labex IBEID, the INCEPTION program, and the Institut Pasteur. We gratefully acknowledge the UTechS Photonic BioImaging (Imagopole), C2RT, Institut Pasteur, supported by the French National Research Agency (France BioImaging, ANR-24-INBS-0005 FBI (BIOGEN); Investments for the Future), and acknowledge support from Institut Pasteur for the use of the Opera Phenix Plus microscope. We thank Y. Vitrenko, L. Lemée and the Biomics Platform (C2RT) at Institut Pasteur (supported by France Génomique (ANR-10-INBS-09) and IBISA) for the generation of RNA sequencing data.

## Supplementary Figure legends

**Figure S1:**
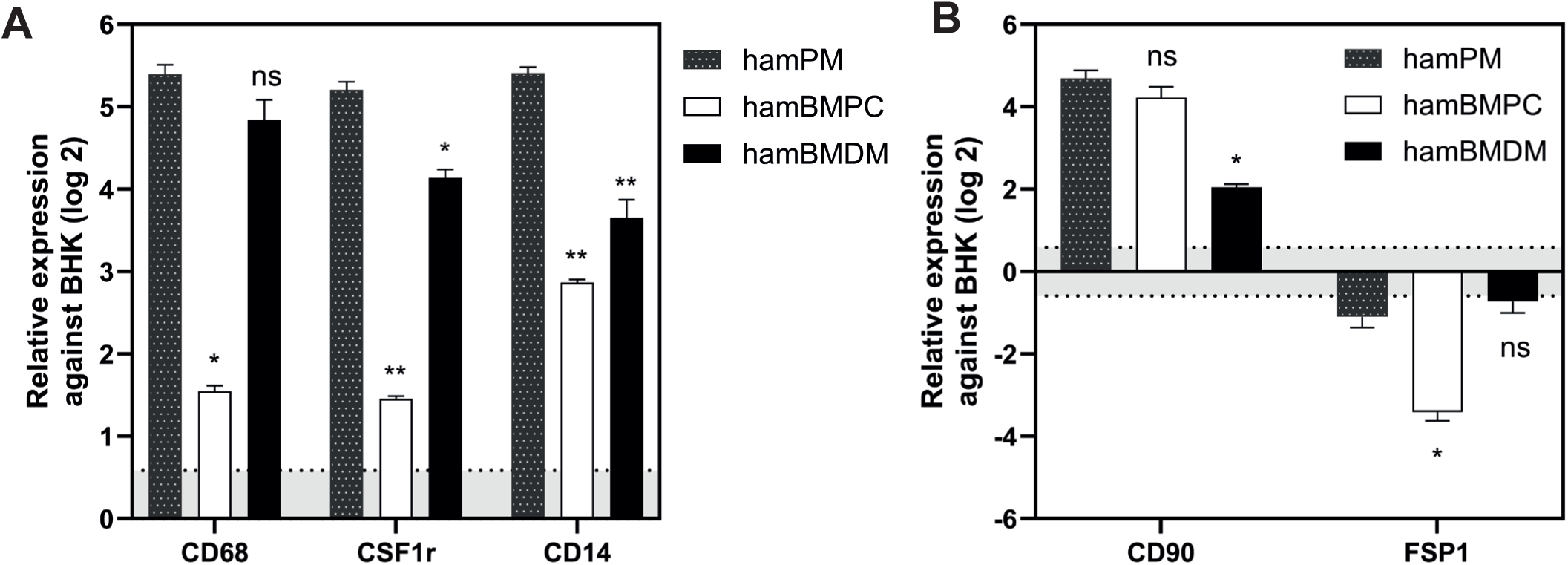
(A-B) RT-qPCR analysis of relative expression of macrophage (A) and fibroblast (B) markers. Total RNA was extracted from hamster progenitor cells (hamBMPC), peritoneal macrophages (hamPM), a hamster fibroblast cell line (BHK), and hamBMDMs after 6 days of differentiation with different sources of CSF1. Dotted black, hamPM; black, rhCSF1-differentiated hamBMDM; white, hamBMPC. Results were calibrated against transcript levels in BHK cells used as positive control for fibroblast expression. Results are expressed as the mean ± SEM of 3 independent experiments. Statistical significance was evaluated using the Tukey’s multiple comparisons, and p-values were calculated for each condition compared to hamPM. Ns, non-significant; *p<0.05; **p < 0.005.

**Figure S2:**
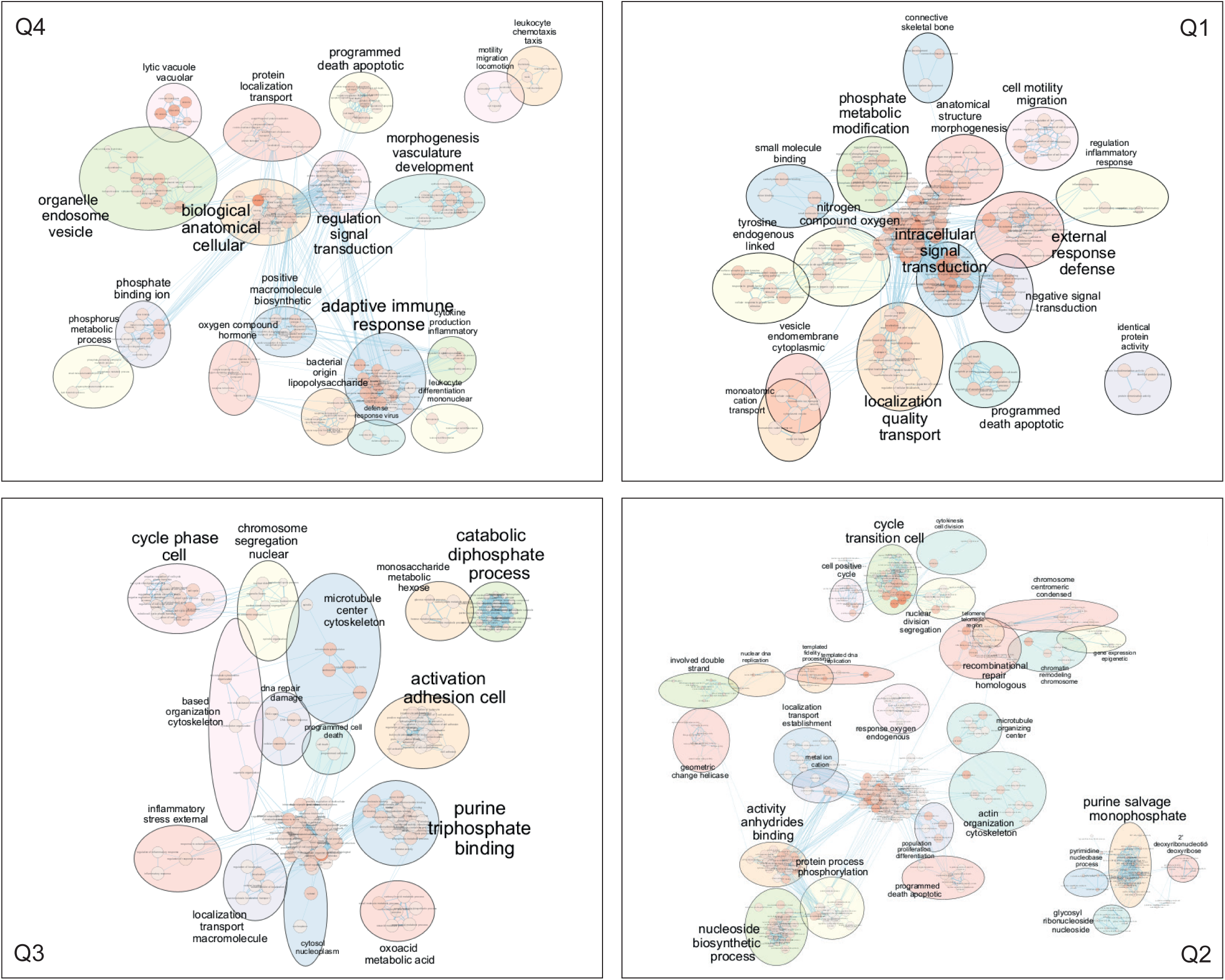
(A) Enrichment maps for each quadrant of the double ratio plot shown in Figure 3A were generated with Cytoscape 3.10.2 using g:Profiler output with a node cutoff 0.05, edge cutoff 0.5 and data set edges – automatic sparse 1/5. Terms and bubble were manually curated and positioned for clarity.

**Figure S3:**
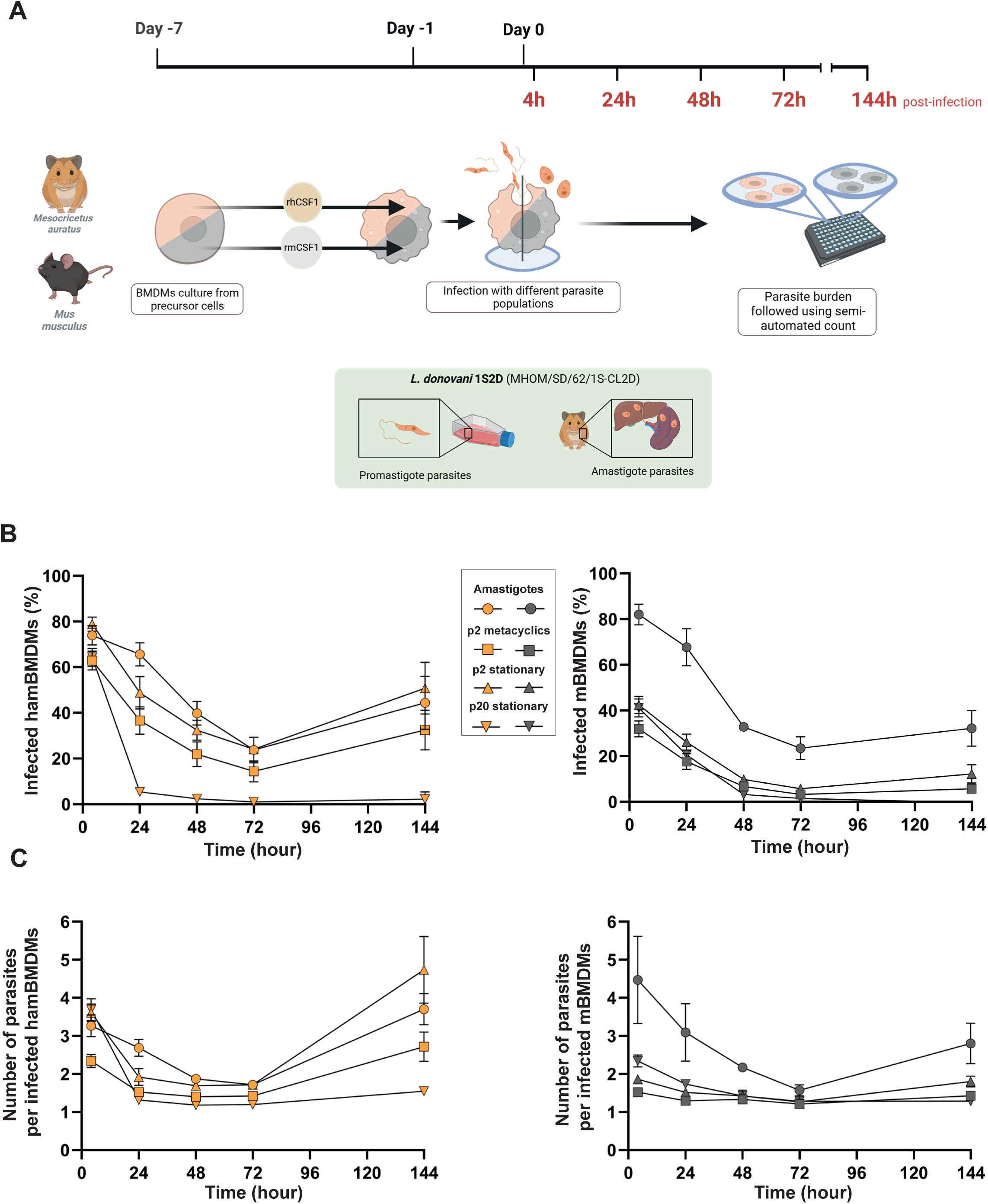
(A) Overview of the experimental workflow for the infection of ham- and mBMDMs with *L. donovani* promastigote and amastigote parasites. (B, C) Evaluation of ham- and mBMDM parasite burden over time after infection with promastigotes and amastigotes according to the legend shown in the graph. The percentage of infected cells (B) and the number of intracellular parasites per macrophage (C) for hamBMDMs (left panels) and mBMDMs (right panels) is reported as a function of time (x-axis).

**Figure S4:**
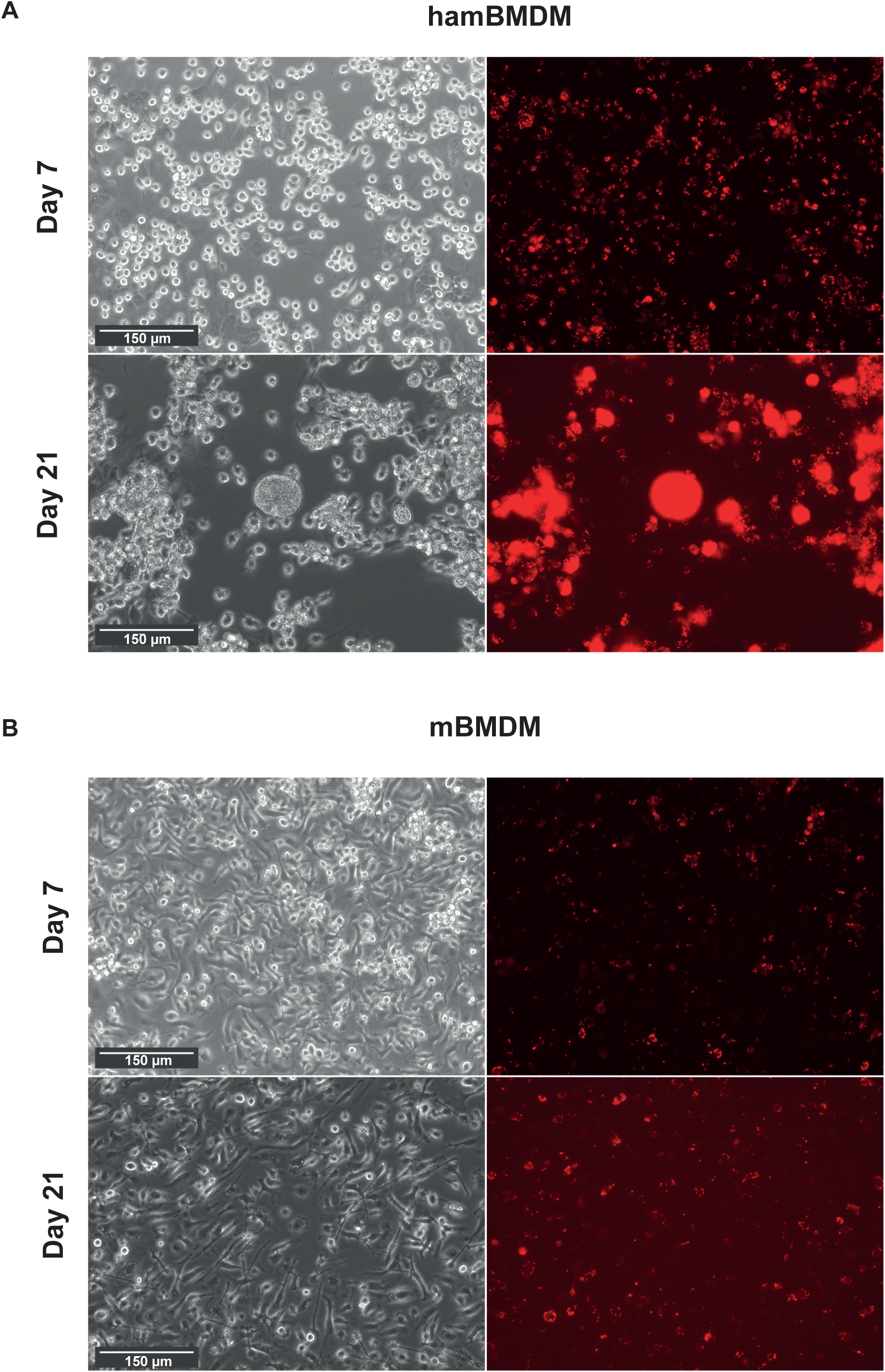
Representative microscopic images of infected hamBMDMs (A) and mBMDMs (B) showing both brightfield (left panels) and parasite *mCherry* fluorescence (right panels) at 7 days (upper panels) and 21 days (lower panels) post-infection.

**Figure S5:**
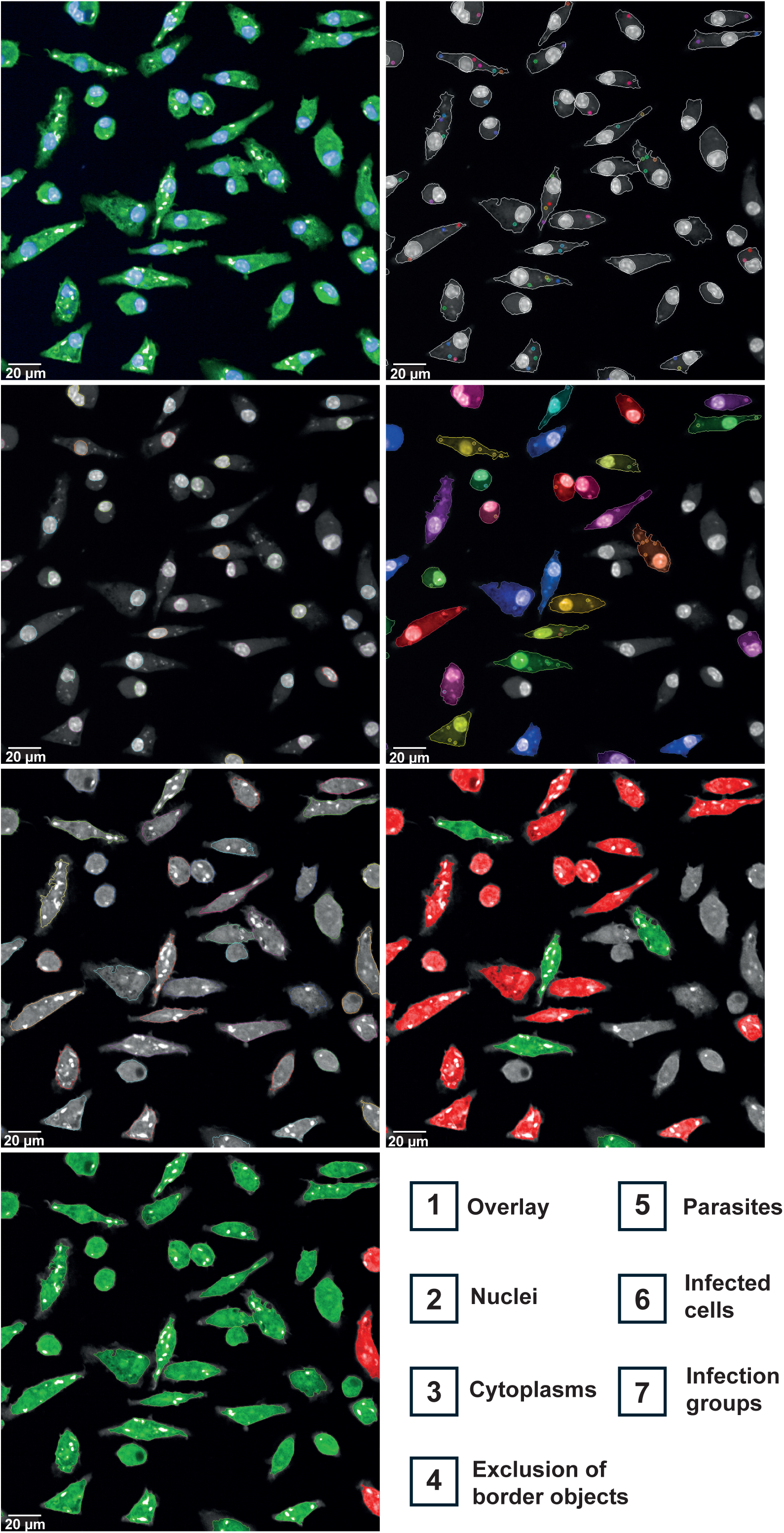
Microscopic images were acquired using an OPERA Phenix Plus confocal microscope and processed applying a custom script developed in collaboration with the BioImagerie Photonique platform (UTechS PBI) at Institut Pasteur, implemented in the SImA software package. The content of each image is indicated in the legend and detailed in the Materials and Methods section.

