## supplemental table 6 for "Intrinsic differences in hamster and mouse macrophage biology correlate with susceptibility to *L. donovani* infection"

|  | **Input** | **Method** | **Output** |
| --- | --- | --- | --- |
| Find Nuclei | Channel : Hoechst 33342 | Common Threshold : 0.5  Area : > 35 µm²  Splitting Coefficient : 3.9  Individual Threshold : 0.33  Contrast : > 0.2 | Nucleus |
| Find Cytoplasm | Channel : Alexa 488 Population: Nuclei | Common Threshold : 0.18  Individual Threshold : 0.15 | Cytoplasm |
| Remove cells at the borders | Population :  Nuclei and cytoplasm | Common Filters  Remove Border Objects | Cells Selected |
| Define macrophage nucleus | Population : Cells Selected Region : Nucleus | Standard  Area  Roundness |  |
| Select nucleus based on channel intensity | Channel : Hoechst 33342  Population : Cells Selected  Region : Nucleus | Standard  Mean | Intensity Nucleus Hoechst 33342 |
| Select macrophage cells | Population : Cells Selected | Filter by Property  Nucleus Area [µm²] : ≤ 200 | Real cells (macrophages) |
| Select fibroblast cells | Population : Cells Selected | Filter by Property  Nucleus Area [µm²] : > 200 | \| Contaminating cells (fibroblasts) |
| Find parasites in macrophages | Channel : Hoechst 33342  Population: Real Cells  Region of Interest (ROI) : Cytoplasm | Radius : ≤ 4 px  Contrast : > 0.32  Uncorrected Spot to Region Intensity : > 1  Distance : ≥ 3 px  Spot Peak Radius : 0 px  Calculate Spot Properties | Spots in  Real Cells |
| Find parasites in fibroblasts | Channel : Hoechst 33342  Population : Contaminating Cells  ROI : Cytoplasm | Radius : ≤ 4 px  Contrast : > 0.4  Uncorrected Spot to Region Intensity : > 1  Distance : ≥ 3 px  Spot Peak Radius : 0 px  Calculate Spot Properties | \| Spots in Contaminating Cells |
| Select infected macrophages | Population : Real Cells | Filter by Property  Number of Spots : > 0 | Infected Real Cells |
